# Nisin- and ripcin-derived hybrid lanthipeptides display selective antimicrobial activity against *Staphylococcus aureus*

**DOI:** 10.1101/2021.04.13.439647

**Authors:** Xinghong Zhao, Oscar P. Kuipers

## Abstract

Lanthipeptides are (methyl)lanthionine ring-containing ribosomally synthesized and post-translationally modified peptides (RiPPs). Many lanthipeptides show strong antimicrobial activity against bacterial pathogens, including antibiotic-resistant bacterial pathogens. The group of disulfide bond-containing antimicrobial peptides (AMPs) is well known in nature and forms a rich source of templates for the production of novel peptides with corresponding (methyl)lanthionine analogues instead of disulfides. Here, we show that novel macrocyclic lanthipeptides (termed thanacin and ripcin) can be synthesized using the known antimicrobials thanatin and rip-thanatin as templates. Notably, the synthesized nisin(1-20)-ripcin hybrid lanthipeptides (ripcin B-G) showed selective antimicrobial activity against *S. aureus*, including an antibiotic-resistant MRSA strain. Interestingly, ripcin B-G, which are hybrid peptides of nisin(1-20) and ripcin, respectively, that are each inactive against Gram-negative pathogens, showed substantial antimicrobial activity against the tested Gram-negative pathogens. Moreover, ripcin B-G was highly resistant against the nisin resistance protein (NSR; a protease could cleave nisin and strongly reduce its activity), opposed to nisin itself. Mode of action studies show that ripcin C exerts its antimicrobial activity against Gram-positive pathogens by binding to the cell wall synthesis precursor lipid II and thereafter arrests cell growth. In addition, ripcin C exerts its antimicrobial activity against Gram-negative pathogens by binding to LPS and the cell wall synthesis precursor lipid II. This study provides an example of converting disulfide bond-based AMPs into (methyl)lanthionine-based macrocyclic hybrid lanthipeptides and can yield antimicrobial peptides with selective antimicrobial activity against *S. aureus*.

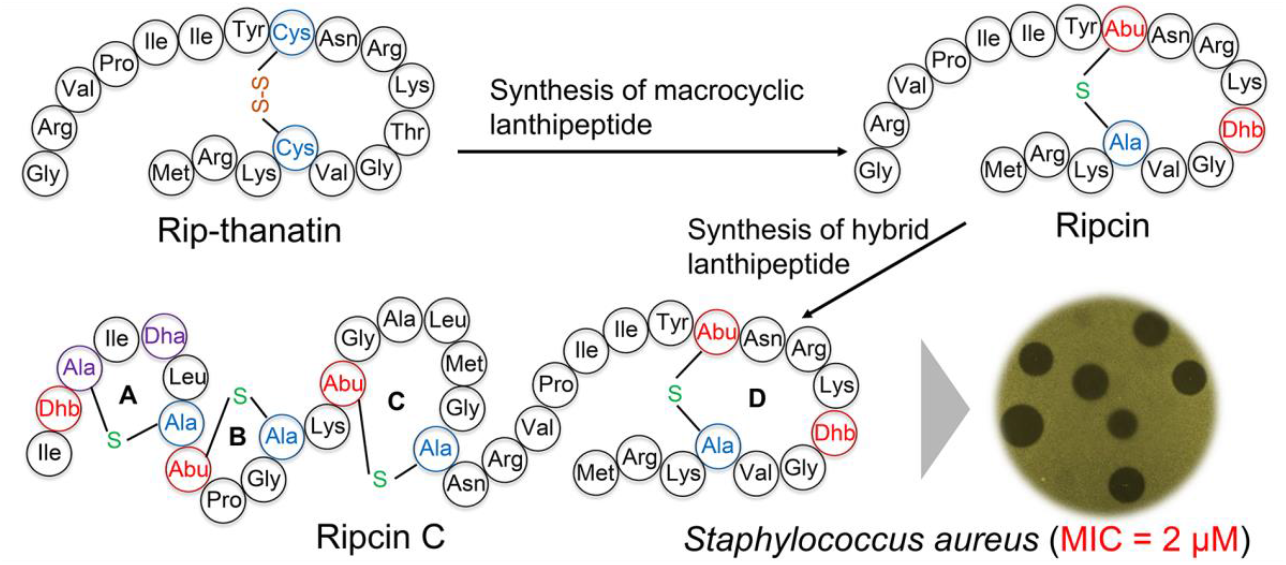
For Table of Contents Use Only

## INTRODUCTION

Lanthipeptides are (methyl)lanthionine ring-containing ribosomally synthesized and post-translationally modified peptides (RiPPs) ^1^. Many lanthipeptides show potent antimicrobial activity against pathogens and/or even against antibiotic-resistant pathogens ^2–7^. Notably, several lanthipeptides, including duramycin, NVB-302, mutacin 1140, and NAI-107 ^3–6,8,9^, have been tested in the clinic or are very close to the start of clinical trials. These all have been demonstrated to display potent antimicrobial activity *in vivo* ^8–11^. The ribosomal synthesis and low substrate specificity of some of the lanthipeptide modification enzymes provide an opportunity to engineer large numbers of novel antimicrobials ^12^.

Nisin, the best-studied lantibiotic, is a 34 amino acid (or 29 amino acids, if one considers a (methyl)lanthionine as a single amino acid) cationic lanthipeptide produced by various *Lactococcus lactis* strains. Because of its potent antimicrobial activity and safety, it has been used as a food preservative for many years. The N-terminal A/B-rings of nisin form a “pyrophosphate cage” that physically interacts with the pyrophosphate of lipid II, resulting in the formation of nisin-lipid II hybrid pores in the target membrane and inhibition of cell wall synthesis via lipid II abduction ^13,14^. Three essential posttranslational modifications of the ribosomally synthesized nisin precursor take place to yield active nisin (Figure 1) ^12,15,16^. NisB is an enzyme that can catalyze the dehydration of Ser and Thr residues in the precursor core peptide. NisC is an enzyme that can catalyze dehydroalanine/dehydrobutyrine with Cys for the formation of (methyl)lanthionine in the precursor core peptide. NisT is a transporter for secreting modified peptides. These modification enzymes consist of the nisin biosynthetic machinery, which has been widely applied to engineer lanthipeptide drug candidates ^2,17–22^.

**Figure 1.**
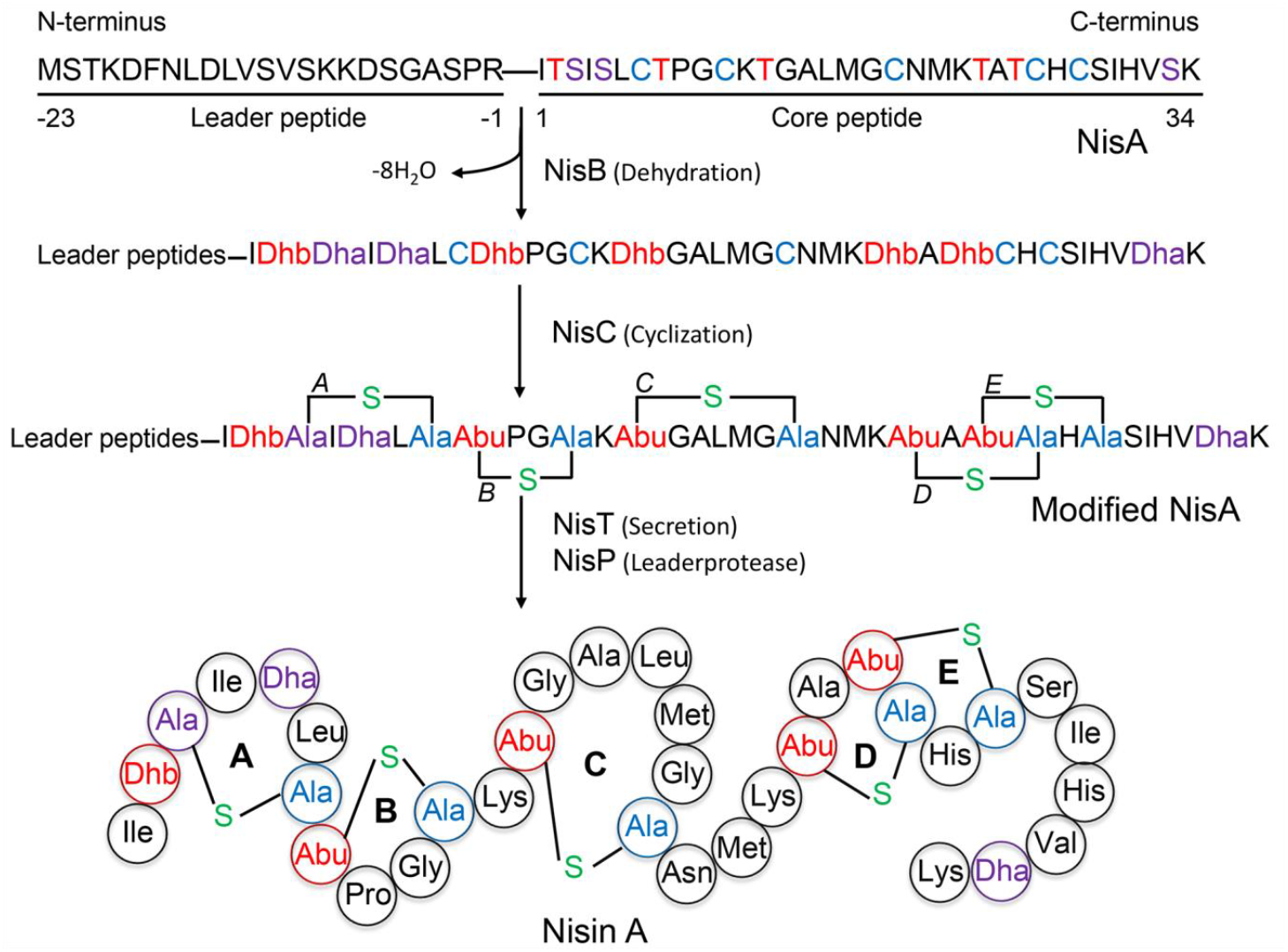
Schematic representation of the biosynthetic route of the model lantibiotic nisin. Dha: dehydroalanine, Dhb: dehydrobutyrine, Abu: aminobutyric acid.

Thanatin (Figure 2A), a 21-residue inducible defense peptide from the hemipteran insect *Podisus maculiventris*, was first reported to exert potent antimicrobial activity against bacteria and fungi in 1996 ^23^. Later studies found that thanatin also has good antimicrobial activity against Gram-negative bacterial pathogens ^24,25^, and mode of action studies show that the antimicrobial activity against Gram-negative bacteria pathogens of thanatin is related to targeting the intermembrane protein complex required for lipopolysaccharide transport ^26^. Rip-thanatin (Figure 2D), an 18-residue insect defense peptide from *Riptortus pedestris*, has also been reported to exert antimicrobial activity against Gram-negative bacteria ^27^. Both rip-thanatin and thanatin are intra-molecular disulfide bond-containing antimicrobial peptides (AMPs). A vast number of such disulfide bond-containing AMPs are known in nature ^28–30^, which form a rich source of templates for producing hybrid peptides with the corresponding (methyl)lanthionine analogs.

**Figure 2.**
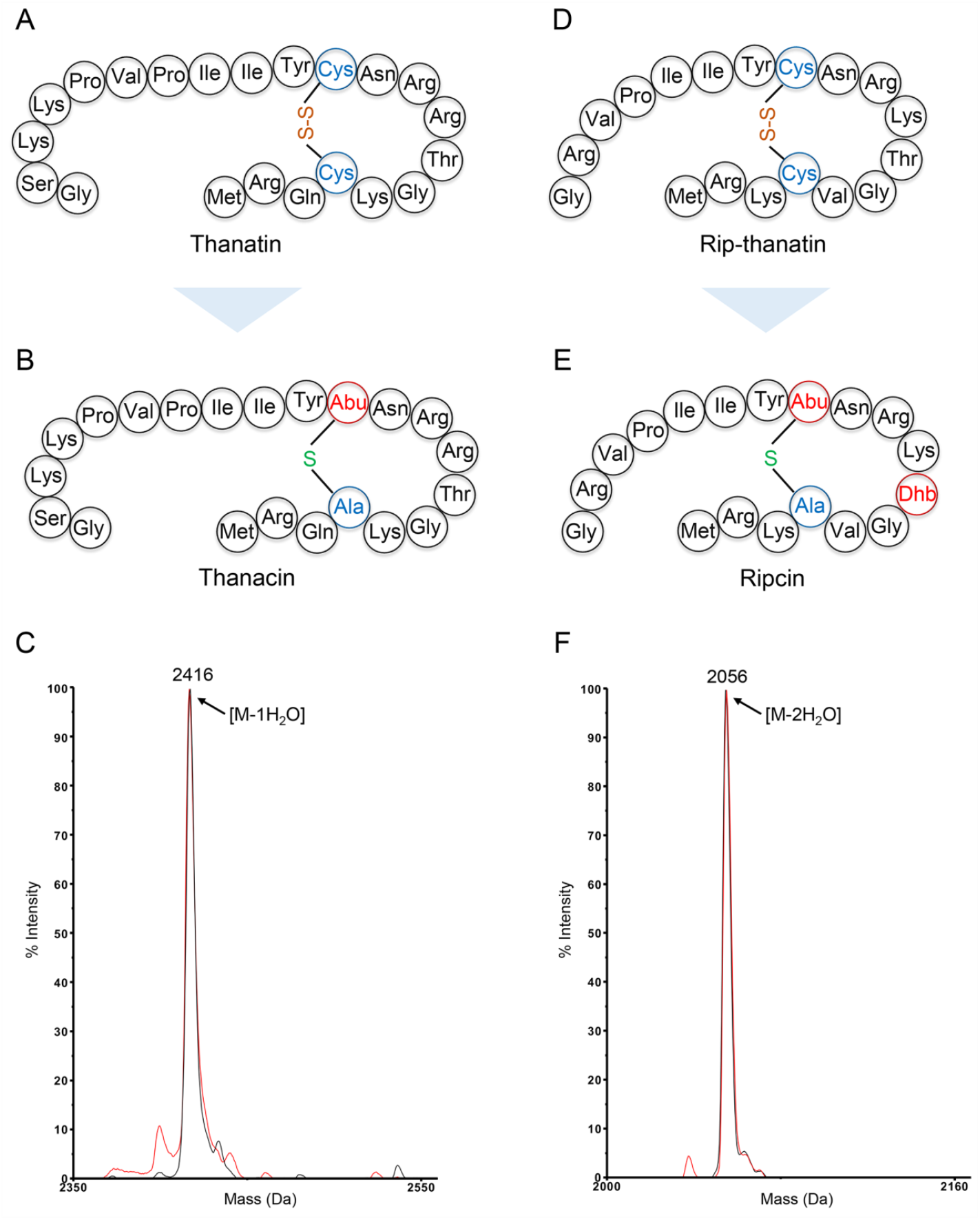
Structures of thanatin (A) and rip-thanatin (D); structures of designed thanacin (B) and ripcin (E); MALDI-TOF MS of thanacin (C) and ripcin (F) before (black) and after (red) CDAP treatment. Dhb: dehydrobutyrine, Abu: aminobutyric acid.

In this study, we describe a strategy in which thanatin and rip-thanatin were used as templates for producing the corresponding (methyl)lanthionine analogs. To this end, the nisin synthetic machinery was used for producing methyllanthionine-stabilized thanatin and rip-thanatin analogs (termed thanacin and ripcin, Figure 2 B and E). A methyllanthionine-stabilized large-ring was successfully introduced into thanacin and ripcin, respectively, which was corroborated by MALDI-TOF MS and LC-MS/MS analysis. However, the first generation of peptides showed insufficient antimicrobial activity against the pathogens tested and showed only antimicrobial activity against *Micrococcus flavus* [Minimal inhibitory concentration (MIC), 4 μM] among the bacterial strains tested. Subsequently, either ripcin or a part of ripcin was genetically fused to the C-terminal end of nisin(1-20) to generate the second generation of 6 macrocyclic peptides that we called ripcin B-G. Ripcin B-G showed stronger antimicrobial activity than either nisin(1-20) or ripcin alone against the Gram-positive pathogens tested. Notably, ripcin B-G showed selective antimicrobial activity against *S. aureus*, including an antibiotic-resistant MRSA strain. Interestingly, the fusion of two inactive peptides, nisin(1-20) and ripcin (or part of ripcin), respectively, also yielded active lanthipeptides against Gram-negative bacterial pathogens. Ripcin C showed the highest antimicrobial activity against the tested Gram-negative and Gram-positive pathogens among all designed peptides. Moreover, ripcin B-G was not sensitive to the nisin resistance protein (NSR; a peptidase that cleaves the C-terminus last 6 amino acids of nisin and strongly reduce its antimicrobial activity), opposed to nisin itself. Mode of action studies showed that ripcin C exerts its antimicrobial activity against Gram-positive pathogens by binding to the cell wall synthesis precursor lipid II and thereafter arrests cell growth. Moreover, ripcin C exerts its antimicrobial activity against Gram-negative pathogens by binding to LPS and the cell wall synthesis precursor lipid II. This study provides an example of converting disulfide bond-based AMPs into hybrid (methyl)lanthionine-based macrocyclic lanthipeptides, and some candidates with selective antimicrobial activity against *S. aureus* (MRSA) were obtained.

## RESULTS AND DISCUSSION

### Synthesis of Macrocyclic Lanthipeptides by Using Thanatin and Rip-thanatin as Templates

To introduce a methyllanthionine-based relatively large ring into thanatin and rip-thanatin ^23,27^, the Cys11 of thanatin and Cys8 of rip-thanatin were designed to be replaced by Thr (Table 1), which can be potentially dehydrated by the NisB dehydratase and subsequently form a methyllanthionine based large ring with Cys18 of thanatin and Cys15 of rip-thanatin by the NisC cyclase. The genes encoding the designed peptides were constructed into a pNZ8048-derived plasmid, a commonly used expression vector for *L. lactis*, next to the *nisA* leader peptide gene (Supplemental Figure S1) ^31^, respectively. After verifying the plasmids by sequencing, *L. lactis* NZ9000 ^32^ with pIL3 BTC ^33^, a plasmid encoding the NisB dehydratase gene, the NisC cyclase gene, and the gene of the transporter NisT (Figure 1), was transformed with these designed plasmids, respectively. After induction and purification, MALDI-TOF MS was used to check the mass of the produced peptides. Ripcin was fully dehydrated as predicted (Table 1, Figure 2F), while only one dehydration was observed for thanacin (Table 1, Figure 2C). We found out this is caused by the substrate specificity of NisB ^34^ and the difference in sequences in ripcin and thanatin: Thr12(KTG) in ripcin is modified, while Thr15(RTG) and Ser2(GSK) in thanacin are not modified. Interestingly, further studies evidenced that the dehydrated amino acid residue in thanacin was the Thr11, the desired position. The formation of the potentially NisC-induced thioether cross-link-based ring was investigated by using 1-cyano-4-dimethylaminopyridinium tetrafluoroborate (CDAP), a compound that reacts with unmodified cysteines in peptides and results in an increase of 25 Da in the peptide’s molecular weight ^7,31,35^. No adduct was observed for either of the designed thanacin or ripcin (Figure 2 C and F), while the mass of a free Cys-containing peptide was shifted entirely with a 25 Da increase (Supplemental Figure S2), indicating that no unmodified cysteines were present in either thanacin or ripcin. These results imply that a thioether cross-link in both thanacin and ripcin were formed. To further characterize the produced thanacin and ripcin molecules, LC-MS/MS analysis was performed. No fragmentation was observed between Thr11 and Cys18 of thanatin and between Thr8 and Cys15 of ripcin (Figure 3 A and B), demonstrating a methyllanthionine-based ring was correctly formed for either thanacin or ripcin (Figure 2 B and E), respectively. Although the nisin biosynthetic machinery has been widely used in lanthipeptide engineering, this is the first time to have been used for the successful synthesis of such macrocyclic lanthipeptides with 6 residues in between the β-methyl-lanthionine forming residues. Many potent antibiotics are macrocyclic peptides, such as polymyxin B, duramycin, daptomycin, and many others ^36–38^. Our results suggest that the synthesized macrocyclic peptides can be used as templates for the development of future antibiotics. For instance, the synthesized macrocyclic lanthipeptides can be used as templates for library construction and screening for therapeutic drug candidates in future studies. High-throughput screening of such libraries has recently been published by us, in collaboration with the Panke group ^18^, showing the feasibility of combining the two approaches.

**TABLE 1.**
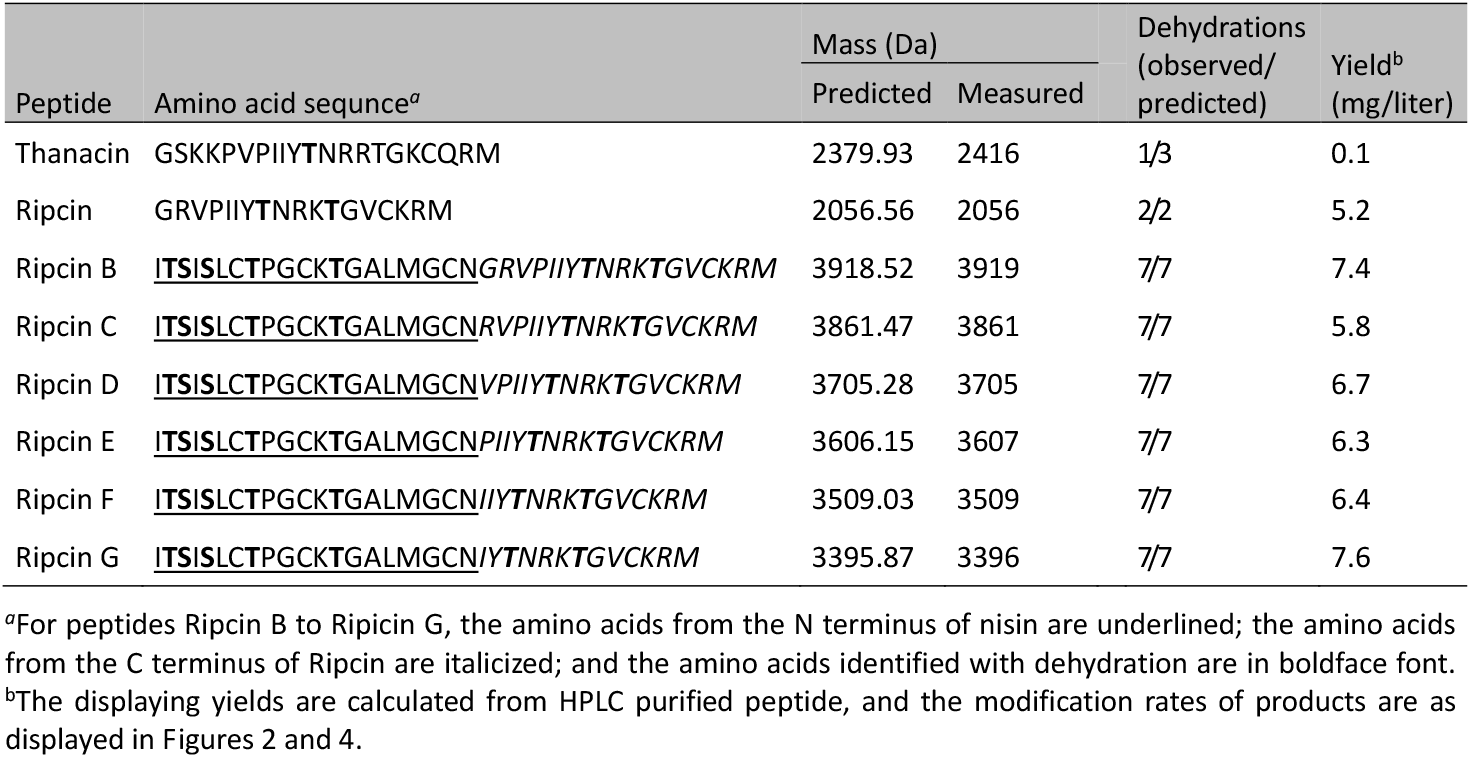
Amino acid sequence, dehydrations and yield of designed peptides

**Figure 3.**
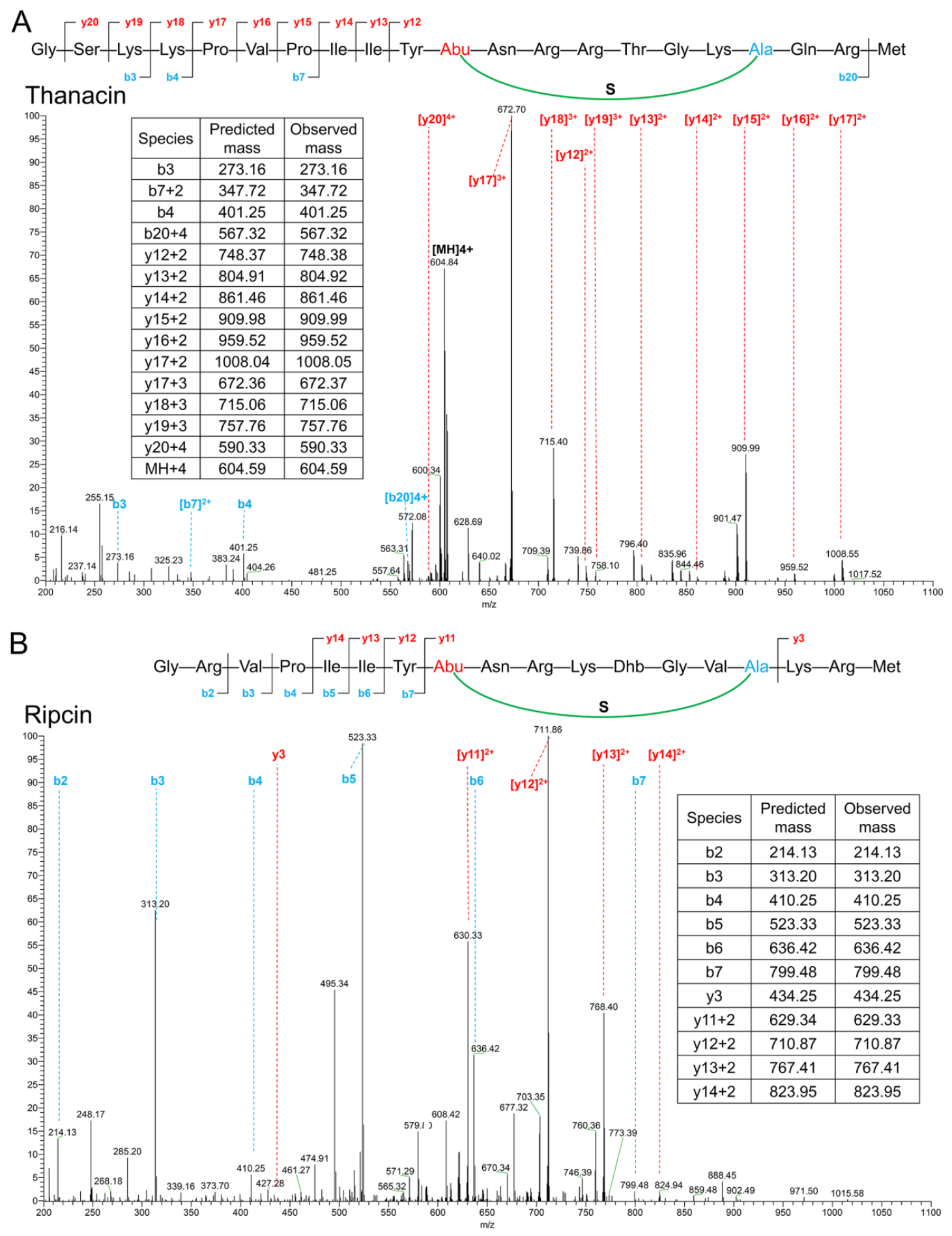
LC-MS/MS spectrum and the proposed structures of thanacin and ripcin. Fragment ions are indicated. (also see Supplemental Figure S3)

### Synthesis of Nisin- and Ripcin-derived Hybrid Lanthipeptides

After HPLC purification, the antimicrobial activities of modified thanacin and ripcin were determined by MIC assays. The results show that both thanacin and ripcin had substantial antimicrobial activity against *M. flavus* (MIC, 4 μM), while both of them had insufficient antimicrobial activity against bacterial pathogens (Table 2, data not shown for thanacin). As ripcin is a large ring- and many positively charged amino acids-containing cyclic peptide and thanacin had a much lower yield than ripcin (Table 1, Supplemental S6), we chose ripcin for further studies. We reasoned that a nisin lipid II binding moiety together with a fused ripcin might display potent antimicrobial activity against bacterial pathogens. Considering the fact that nisin(1-20) and ripcin-fused peptide (ripcin B) would contain 38 amino acids, which might be too long to be modified by the NisB dehydratase and NisC cyclase, five shorter peptides (Ripcin C-G) were designed by using nisin(1-20) and a part of ripcin (Table 1), respectively. The genes encoding the designed peptides were constructed into a pNZ8048-derived plasmid next to the *nisA* leader peptide gene (Supplemental Figure S4 and S5) ^31^, respectively. After verification of the plasmids by sequencing, *L. lactis* NZ9000 ^32^ with pIL3 BTC ^33^ was transformed with these plasmids, respectively. After induction and purification, MALDI-TOF MS was used to measure the mass of the produced peptides. All of the six designed peptides were dehydrated as predicted (Table 1, Figure 4), and the yields of the designed peptides were between 5.8 mg/L to 7.6 mg/L (Table 1, Supplemental S6), which is a relatively high yield compared to previously reported nisin-derived peptides ^39^. A CDAP assay showed that no adduct was observed for all of the six designed peptides (Figure 4), indicating the peptides were modified as predicted (Figure 5). In a previous study, the nisin synthetic machinery was successfully applied to modify a peptide with 9 dehydrations and 7 rings ^7^. Van Heel et al ^19^ reported that flavucin, which was discovered by genome mining, was successfully modified by the nisin synthetic machinery. Overall, these studies on the production and secretion of modified peptides, show that the nisin synthetic machinery provides an efficient lanthipeptide engineering system.

**TABLE 2.** Antimicrobial activity of designed peptides against microorganisms

| Organism and type<sup>a</sup> | MIC (μM) <sup>b</sup> |  |  |  |  |  |  |  |  |  |
| --- | --- | --- | --- | --- | --- | --- | --- | --- | --- | --- |
|  | Nisin(1-20) | Ripcin | Ripcin B | Ripcin C | Ripcin D | Ripcin E | Ripcin F | Ripcin G | Nisin | Polymyxin B |
| **Gram-positive bacteria** |  |  |  |  |  |  |  |  |  |  |
| *Staphylococcus aureus* ATCC15975 (MRSA) | 32 | > 256 | 4 | 2 | 8 | 4 | 4 | 4 | 2 | 32 |
| *Staphylococcus aureus* LMG01041 | 32 | > 256 | 2 | 2 | 4 | 4 | 4 | 4 | 4 | 32 |
| *Enterococcus faecium* LMG16003 (VRE) | 64 | > 256 | 16 | 8 | 32 | 32 | 16 | 16 | 2 | 16 |
| *Enterococcus faecalis* LMG16216 (VRE) | 128 | > 256 | 32 | 16 | 32 | 32 | 32 | 32 | 2 | 16 |
| *Bacillus cereus* ATCC 14579 | 64 | > 256 | 32 | 16 | 16 | 16 | 16 | 32 | 2 | 8 |
| *Lactococcus lactis* NZ9000 (pNZ8048-Em<sup>r</sup>) | 4 | > 256 | 0.5 | 0.25 | 0.5 | 0.5 | 0.5 | 0.5 | **0.02** | > 32 |
| *Lactococcus lactis* NZ9000 (pNZ-SV- SaNSR) | 4 | > 256 | 0.5 | 0.25 | 0.5 | 0.5 | 0.5 | 0.5 | **2** | > 32 |
| **Gram-negative bacteria** |  |  |  |  |  |  |  |  |  |  |
| *Acinetobacter baumannii* LMG01041 | > 256 | > 256 | 8 | 4 | 32 | 32 | 16 | 32 | 4 | 1 |
| *Acinetobacter baumannii* LMG17978 | > 256 | > 256 | 64 | 32 | > 128 | > 128 | > 128 | > 128 | 32 | 4 |
| *Shigella flexneri* ATCC29903 | > 256 | > 256 | 16 | 8 | 32 | 32 | 32 | 32 | 16 | 0.5 |
| *Escherichia coli* LMG8223 | > 256 | > 256 | 16 | 16 | 32 | 64 | 32 | 64 | 16 | 2 |
| *Klebsiella pneumoniae* LMG20218 | > 256 | > 256 | 64 | 32 | > 128 | > 128 | > 128 | > 128 | 64 | 2 |
| *Pseudomonas aeruginosa* LMG6395 | > 256 | > 256 | > 128 | 64 | > 128 | > 128 | > 128 | > 128 | 64 | 2 |
| *Enterobacter cloacae* LMG02783 | > 256 | > 256 | 64 | 32 | > 128 | > 128 | > 128 | > 128 | 32 | > 128 |

<sup>a</sup>VRE, vancomycin-resistant enterococci; MRSA, oxacillin-methicillin-resistant *Staphylococcus aureus*.
<sup>b</sup>The MIC was determined by broth microdilution. Nisin and Polymyxin B were used as well-known antibiotic controls.

**Figure 4.**
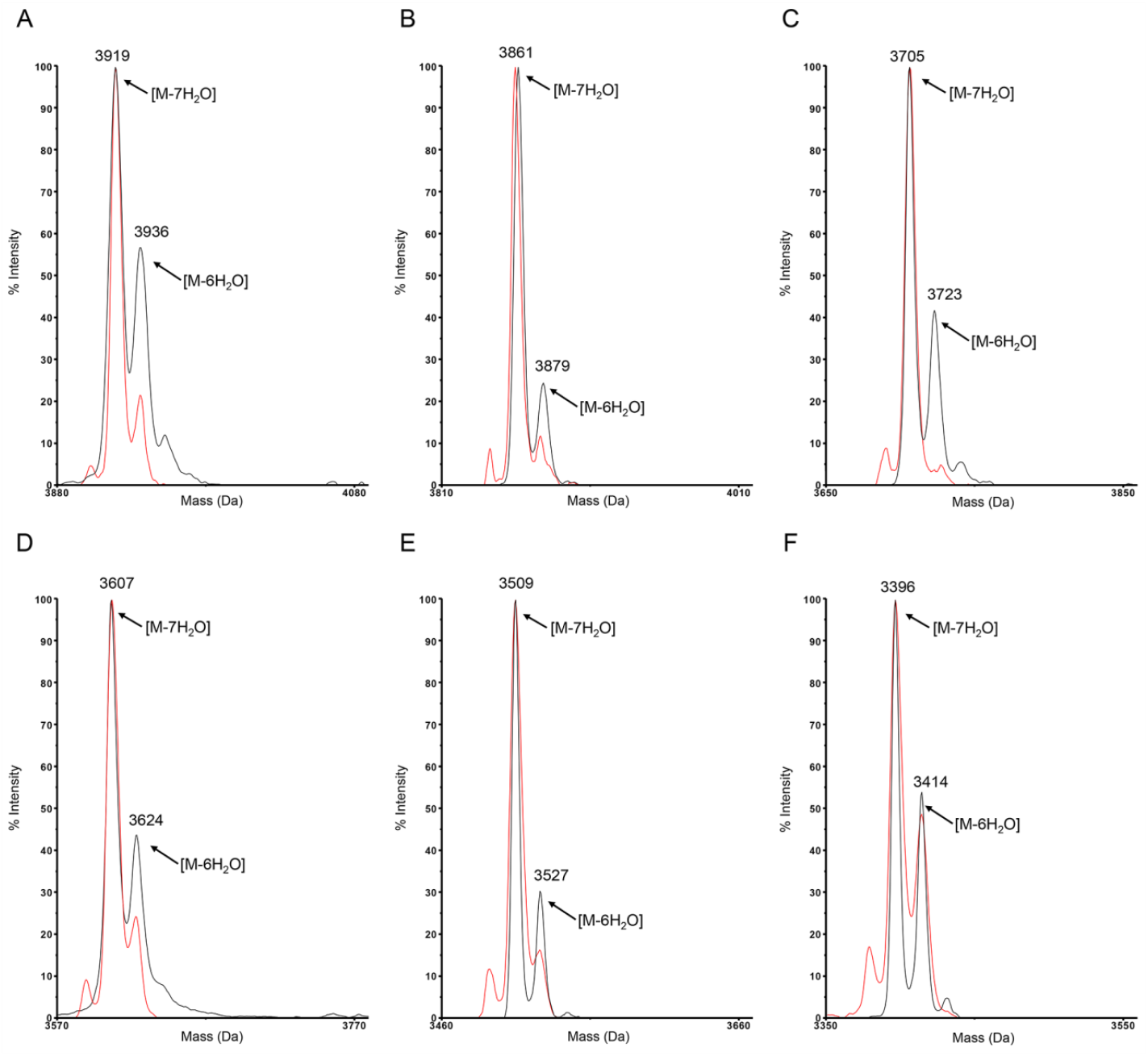
MALDI-TOF MS of ripcins before (black) and after (red) CDAP treatment. **A**, ripcin B; **B**, ripcinC; **C**, ripcin D; **D**, ripcin E; **E**, ripcin F; **F**, ripcin G.

**Figure 5.**
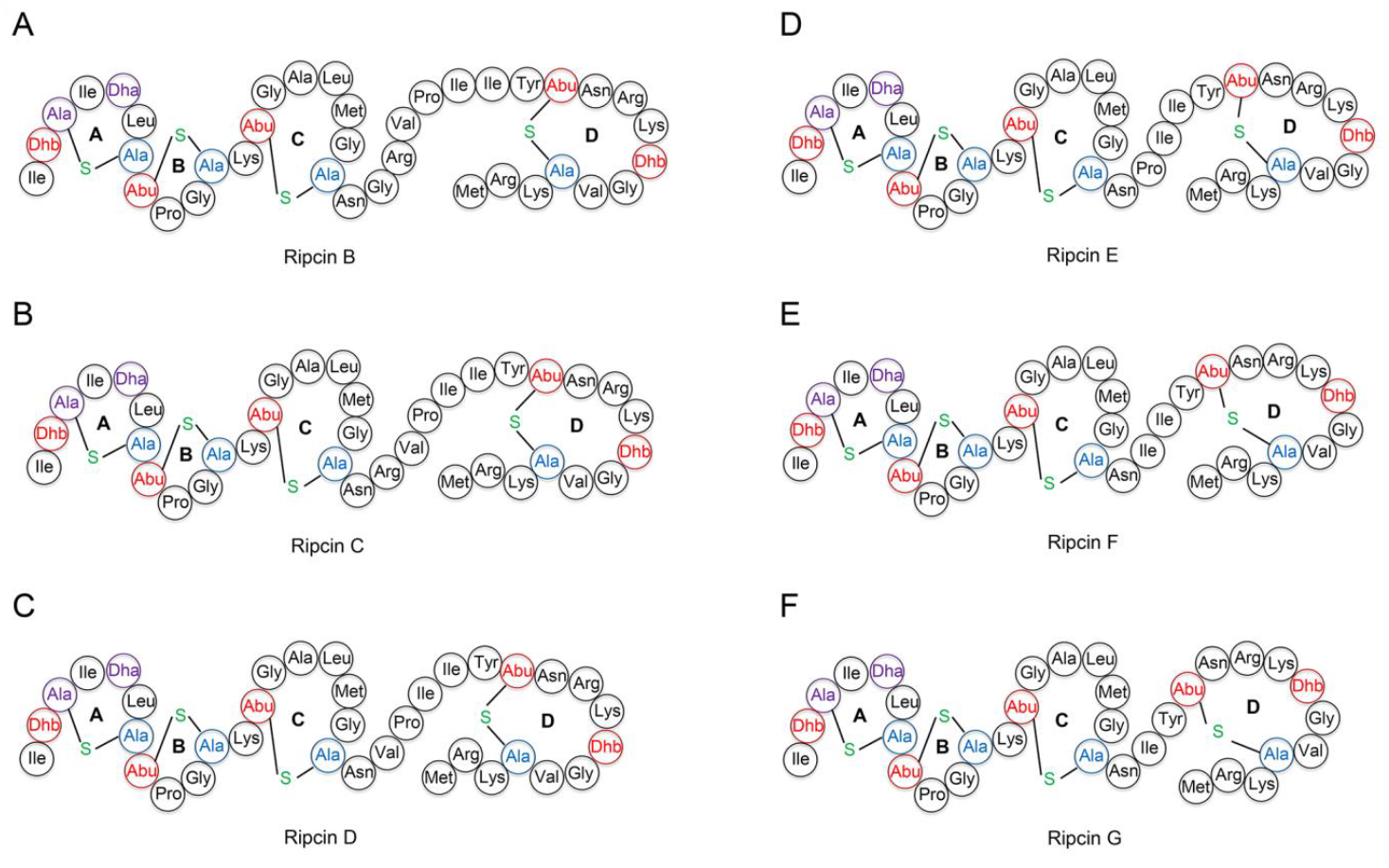
Hypothetical structures of ripcins. **A**, ripcin B; **B**, ripcinC; **C**, ripcin D; **D**, ripcin E; **E**, ripcin F; **F**, ripcin G. Dha: dehydroalanine, Dhb: dehydrobutyrine, Abu: aminobutyric acid.

### Ripcin B-G Show Selective Antimicrobial Activity Against *S. aureus*

To determine the antimicrobial activity of ripcin B-G against bacterial pathogens, a MIC assay was performed according to the standard guidelines ^40^. Nisin and polymyxin B were used as antibiotic controls. Ripcin showed neither antimicrobial activity against the tested Gram-positive nor Gram-negative bacterial pathogens. Nisin(1-20) was obtained using chymotrypsin to digest full nisin (Figure S7 and S8) ^41^. Nisin(1-20) showed insufficient antimicrobial activity against all tested Gram-positive pathogens, while it showed no antimicrobial activity against the tested Gram-negative pathogens (Table 2). Ripcin B-G showed stronger antimicrobial activity against the tested Gram-positive pathogens than nisin(1-20). Notably, ripcin B-G showed selective antimicrobial activity against *S. aureus* (Table 2), including an antibiotic-resistant MRSA strain. The antimicrobial activity of ripcin B-G against *S. aureus* is comparable with that of the well-known antimicrobial RiPP nisin.

Many studies have shown that recombination of antimicrobial peptides forms an alternative strategy for the synthesis of antibiotics with specific antimicrobial activity ^7,42,43^. Taken together, these studies and our results suggest that additional hybrid antimicrobials can be synthesized for developing antibiotics with a specific antimicrobial activity. Interestingly, ripcin B-G, which are recombinant peptides of two inactive (against Gram-negative pathogens) peptides nisin(1-20) and ripcin, showed substantial antimicrobial activity against the tested Gram-negative bacteria (Table 2). Consistently, a previous study showed that recombination of the nisin N-terminus part with cationic antimicrobial peptides could increase the antimicrobial activity of nisin against Gram-negative pathogens ^39^.

To investigate the stability of ripcin B-G against the nisin resistance protein (NSR) ^44,45^, a peptidase that cleaves the C-terminus last 6 amino acids of nisin, a MIC test was performed by using NSR producing strain *L. lactis* NZ9000 (pNZ-SV-SaNSR, Em^r^) ^45^ and non-NSR producing strain *L. lactis* NZ9000 (pNZ8048-Em^r^, Em^r^) ^46^. Due to cleave of the C-terminus last 6 amino acids of nisin by NSR protease, the antimicrobial activity of nisin against *L. lactis* NZ9000 decreased 100-fold in the presence of the NSR protease (Table 2). Interestingly, ripcin B-G, respectively, showed no reduction of antimicrobial activity against *L. lactis* NZ9000 in the presence of the NSR protease (Table 2). These results demonstrate that the hybrid macrocyclic lanthipeptides, ripcin B-G, bypassed the NSR resistance mechanism towards nisin. Together, the here-synthesized lanthipeptides with a large C-terminal ring, named ripcin B-G, have shown the potential to be developed as antibiotics for treatment of *S. aureus*-caused infections without the concern of NSR. Among the designed peptides, ripcin C showed the highest antimicrobial activity against the tested bacterial pathogens. Therefore, ripcin C was selected for further studies.

### Ripcin C Acts as a Bacteriostatic Antibiotic Against Gram-positive Pathogens, While it Shows Bactericidal Activity Against Gram-negative Pathogens

To investigate whether Ripcin C acts as a bacteriostatic or bactericidal antibiotic against bacteria pathogens, time-killing assays were performed. *S. aureus* (MRSA) and *A. baumannii* LMG01041 were inoculated in MHB and grown until the OD_600_ of cell cultures reached 0.8. Cell cultures were then diluted to a concentration of 5×10^6^ c.f.u. per mL and thereafter challenged with antibiotics at a concentration of 10×MIC. Nisin was used as a bactericidal antibiotic control against Gram-positive bacteria ^13^. Nisin killed all of the *S. aureus* (MRSA) in 2 h, while ripcin C did not reduce the population of *S. aureus (MRSA)* during 8 h after treatment (Figure 6A). These results demonstrate that ripcin C acts as a bacteriostatic antibiotic against Gram-positive bacteria. These results are in line with the later membrane permeability assay and lipid II binding assay, which showed that ripcin C binds to lipid II (Lys) but does not disrupt the cell membrane (Figure 6C and 7B). Polymyxin B was used as a bactericidal antibiotic control against Gram-negative bacteria ^47^. Polymyxin B showed a quick killing capacity on *A. baumannii* cells, which killed all bacteria in half an hour (Figure 6B). Ripcin C showed slower bactericidal activity than polymyxin B against *A. baumannii* cells, which killed all Gram-negative bacteria in 2 h (Figure 6B).

**Figure 6.**
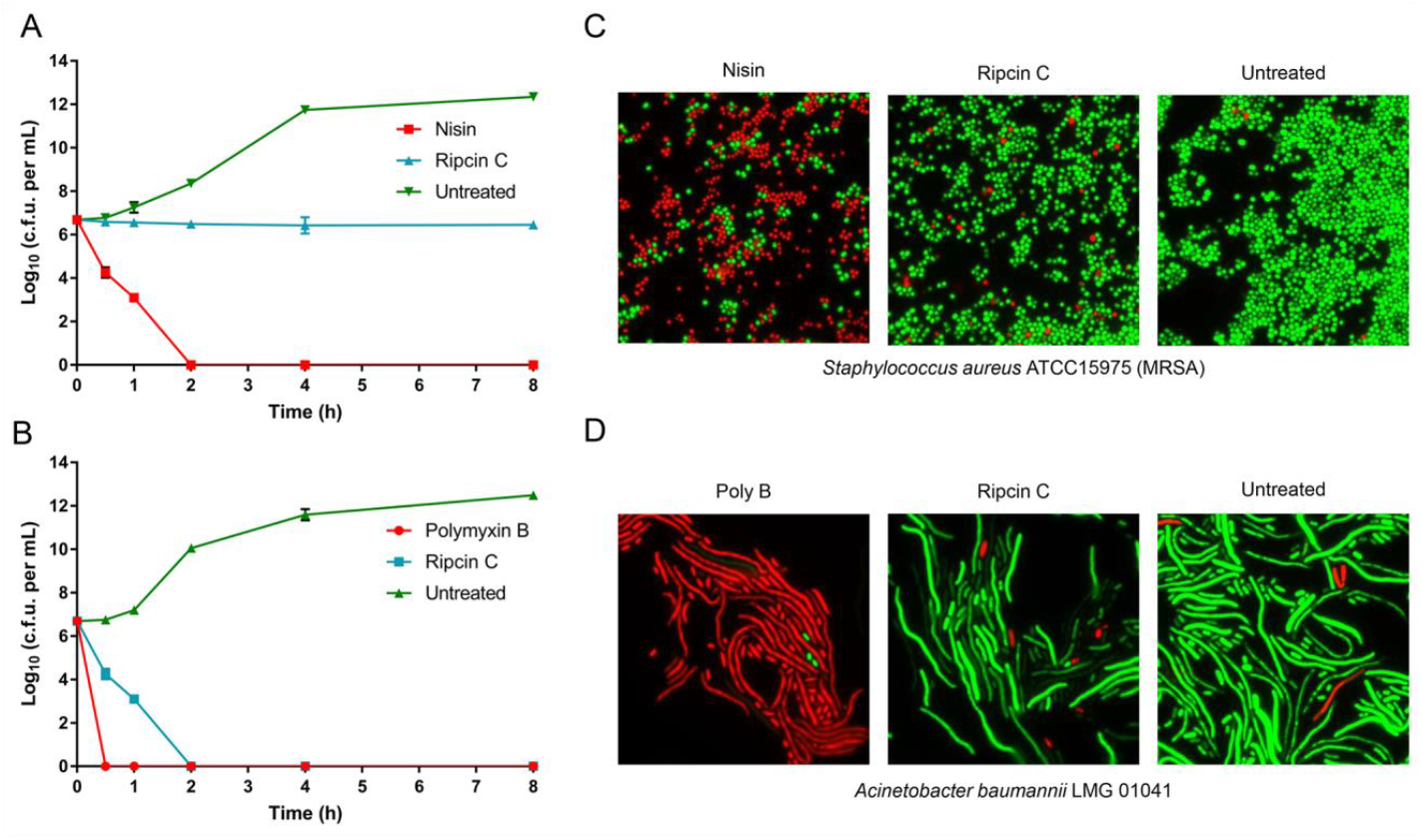
Ripcin C acts as a bacteriostatic antibiotic against Gram-positive pathogens and shows bactericidal activity against Gram-negative pathogens. **A**, time-killing assay of ripcin C, *S. aureus* (MRSA) was challenged with lantibiotics (10×MIC). Data are representative of 3 independent experiments ± the standard deviation (SD). **B**, time-killing assay of ripcin C, *A. baumannii* was challenged with lantibiotics (10×MIC). Data are representative of 3 independent experiments ± the standard deviation (SD). **C**, fluorescence microscopy images of *S. aureus* (MRSA), which was challenged with different antibiotics at a concentration of 2×MIC. **D**, fluorescence microscopy images of *A. baumannii*, which was challenged with different antibiotics at a concentration of 2×MIC. In panels **C** and **D**, green denotes a cell with an intact membrane, whereas red denotes a cell with a compromised membrane.

### Ripcin C does not Disrupt the Membrane of Bacteria

To assess the influence of ripcin C on the bacterial membrane, membrane permeability assays were performed by using a commercial LIVE/DEAD Baclight Bacterial Viability Kit, which contains SYTO^®^9 and propidium iodide. Cells with an intact membrane will stain green, whereas cells with a compromised membrane will stain red. After treatment with ripcin C at a concentration of 2×MIC for 15 min, the cells were monitored by fluorescence microscopy. Green cells were observed for both *S. aureus* (MRSA) and *A. baumannii* (Figure 6 C and D), indicating that ripcin C exerts its antimicrobial activity, not via disruption of the cellular membrane.

### Ripcin C Binds to the Cell Wall Synthesis Precursor Lipid II and LPS

To assess the functionality of the lipid II binding motive of ripcin C, bacterial growth assays were performed with or without purified lipid II (Figure 7A) in the presence of ripcin C at a concentration of 2×MIC. For the Gram-positive lipid II (Lys) binding assay, nisin and ampicillin were used as lipid II binding and non-lipid II binding antibiotic controls. Ripcin C inhibited the growth of *S. aureus* (MRSA) cells, while *S. aureus* (MRSA) cells treated with ripcin C were grown in the presence of 10 μM Gram-positive lipid II (Lys) (Figure 7B), indicating ripcin C still keeps the lipid II binding capacity of nisin. For Gram-negative lipid II (Dap) binding assay, nisin and amikacin were used as lipid II binding and non-lipid II binding antibiotic controls, respectively. In the presence of 10 μM Gram-negative lipid II (Dap), ripcin C lost its antimicrobial activity against *A. baumannii* LMG01041 cells (Figure 7C), indicating ripcin C also binds with the Gram-negative lipid II (Dap). For the lipopolysaccharide (LPS) binding assay, polymyxin B and amikacin were used as LPS binding and non-LPS binding antibiotic controls. Interestingly, in the presence of 100 μg/mL LPS, ripcin C lost its antimicrobial activity against *A. baumannii* cells (Figure 7D), indicating ripcin C has an LPS binding capacity. The LPS binding capacity of ripcin C maybe one of the primary reasons that ripcin C exerts its antimicrobial activity against Gram-negative pathogens, since nisin(1-20) lacks antimicrobial activity against Gram-negative pathogens. In addition, the C-terminus of ripcin C may target the intermembrane protein complex required for lipopolysaccharide transport as thanatin does ^26^, which is probably the reason it shows bactericidal activity against Gram-negative bacteria since binding to lipid II can only arrest bacterial growth. These results provide a primary data for the mode of action of the synthesized hybrid lanthipeptides.

**Figure 7.**
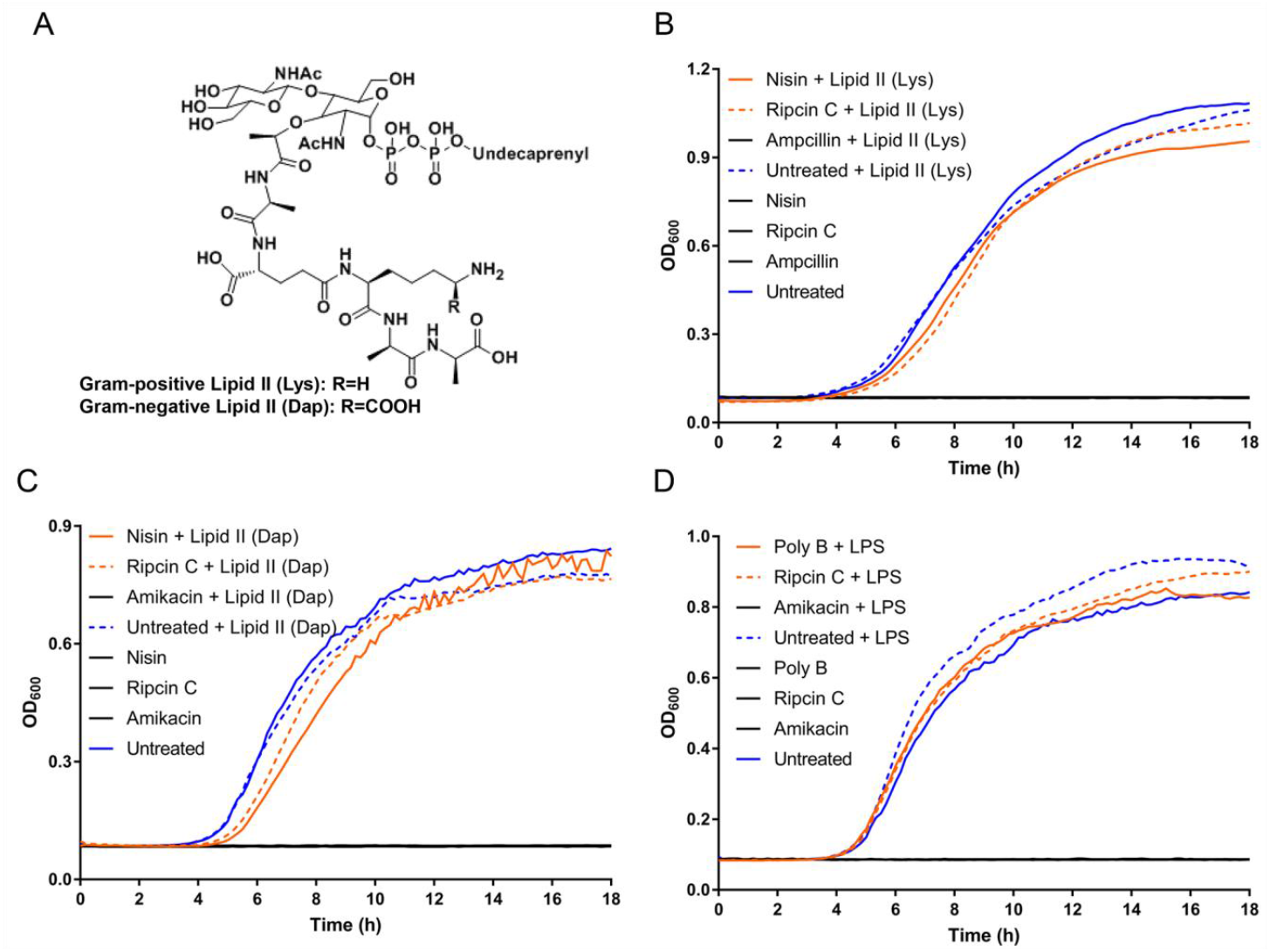
Ripcin C binds with the cell wall synthesis precursor Lipid II and LPS. **A**, structure of Gram-positive Lipid II (Lys) and Gram-negative Lipid II (Dap). **B**, addition of purified Lipid II (Lys) (10μM) inhibited the antimicrobial activity of Ripcin C (2×MIC) against *Staphylococcus aureus* ATCC15975 (MRSA), indicating the binding of Ripcin C and Lipid II (Lys). **C**, addition of purified Lipid II (Dap) (10μM) inhibited the antimicrobial activity of Ripcin C (2×MIC) against *Acinetobacter baumannii* LMG01041, indicating the binding of Ripcin C and LPS. **D**, addition of purified LPS (100μg/mL) inhibited the antimicrobial activity of Ripcin C (2×MIC) against *Acinetobacter baumannii* LMG01041, indicating the binding of Ripcin C and Lipid II (Dap). In panels **B**, **C**, and **D**, all growth curves without bacteria grown are shown in a black solid line.

## CONCLUSIONS

Here, we show that macrocyclic lanthipeptides (thanacin and ripcin) can be synthesized by using thanatin and rip-thanatin as templates. This is the first time that the nisin synthetic machinery was used successfully to synthesize such a macrocyclic lanthipeptide (6 residues within the ring). The synthesized macrocyclic lanthipeptides can be used as templates for library construction and screening for therapeutic drug candidates in future studies. In addition, nisin(1-20) and ripcin hybrid lanthipeptides showed stronger antimicrobial activity than either nisin(1-20) or ripcin alone against the tested bacterial pathogens. Notably, ripcin B-G showed selective antimicrobial activity against *S. aureus*, including an antibiotic-resistant MRSA strain. Interestingly, ripcin B-G, which are hybrid peptides of two inactive (against Gram-negative pathogens) peptides, i.e., nisin(1-20) and ripcin, showed substantial antimicrobial activity against the tested Gram-negative pathogens. Moreover, ripcin B-G were fully resistant against the NSR, while efficient degradation takes place with full nisin ^44,45^, making our hybrid peptides more attractive for application in complex microbial environments like the gut or skin, also in view of their higher target specificity. Mode of action studies show that ripcin C exerts its antimicrobial activity against Gram-positive pathogens by binding to the cell wall synthesis precursor lipid II and thereafter arrests cell growth. Moreover, ripcin C exerts its antimicrobial activity against Gram-negative pathogens by binding to LPS and the cell wall synthesis precursor lipid II. Together, this study shows a convenient approach for converting disulfide bond-based AMPs into (methyl)lanthionine-based macrocyclic lanthipeptides, yielding hybrid macrocyclic lanthipeptides with selective antimicrobial activity against *S. aureus*.

## MATERIALS AND METHODS

### Microbial Strains Used and Growth Conditions

Strains and plasmids used in the present study are listed in Supplemental Table S1. *L. lactis* NZ9000 was used for plasmid construction, plasmid maintenance, and peptide expression. For plasmid selection, *L. lactis* was grown at 30 °C in M17 broth (Oxoid) or M17 broth solidified with 1% (wt/vol) agar, containing 0.5% (wt/vol) glucose (GM17), when necessary, supplemented with chloramphenicol (5 μg/ml) and/or erythromycin (5 μg/ml). For protein expression, stationary-phase cultures, which were gown in GM17, were inoculated (20-fold diluted) on minimal expression medium (MEM) and induced with nisin (5 ng/ml) at an optical density at 600 nm (OD_600_) of about 0.35.

### Molecular Biology Techniques

Oligonucleotide primers used for cloning and sequencing in this study are listed in Supplemental Table S2, and all the oligonucleotide primers were purchased from Biolegio B.V. (Nijmegen, The Netherlands). Constructs coding for designed peptides were made by amplifying template plasmid using a phosphorylated downstream sense- (or upstream antisense) primer and an upstream antisense (or downstream sense) primer with a peptide-encoding tail. The DNA amplification was carried out by using phusion DNA polymerase (Thermo Fisher Scientific, Waltham, MA). Self-ligation of the resulting plasmid was carried out with T4 DNA ligase (Thermo Fisher Scientific, Waltham, MA). The electrotransformation of *L. lactis* was carried out as previously described using a Bio-Rad gene pulser (Bio-Rad, Richmond, CA) ^48^. The designed peptide mutations were verified by sequencing using the primer PrXZ12 at Macrogen Europe B.V.

### Small Scale Expression, and Trichloroacetic Acid (TCA) Precipitation of Peptides for Tricine-SDS Protein Gel Assay

*L. lactis* NZ9000 cells with pIL3 BTC were transformed with peptide plasmids (50 ng), plated on GM17 agar plates supplemented with chloramphenicol (5 μg/ml) and erythromycin (5 μg/ml) (GM17CmEm) and grown at 30 °C for 20 h. A single colony was used to inoculate 5 mL of GM17CmEm broth. The culture was grown overnight at 30 °C and then used to inoculate 45 mL (20-fold dilution) of MEM. After induction at 30 °C for 3h, the supernatant of cultures were harvested by centrifugation at 10,000 × g for 30 min. Ice-cold 100% TCA was added to the ice-cold supernatants to a final concentration of 10%, and samples were subsequently kept on ice for 2 h to precipitate peptides. Samples were then centrifuged at 10,000 × g at 4 °C for 30 min. The precipitate was washed three times with 20 mL ice-cold acetone to remove any residual TCA. Samples were dried in the fume hood and resuspended in 0.5 mL 0.1 M PBS buffer. Subsequently, peptides were separated by Tricine-SDS gel (16%) electrophoresis and visualized by Coomassie Blue staining.

### Large Scale Expression, and Purification of Designed Peptides

*L. lactis* with pIL3 BTC and the corresponding peptide was inoculated on 50 mL of GM17CmEm. After overnight grown at 30 °C, cultures were inoculated on 1 L (20-fold dilution) of MEM. After induction, cultures were grown at 30 °C for overnight. After centrifugation of the overnight expressed cultures (OD_600_ ≈1.2), the supernatants were collected and adjusted pH to 7.0. After that, the cultures were applied to CM Sephadex^TM^ C-25 column (GE Healthcare) equilibrated with distilled water. The flow-through was discarded, and the column was subsequently washed with 12 column volumes (CV) of distilled water. The peptide was eluted with 6 CV 2 M NaCl. The eluted peptide was then applied to a SIGMA-ALDRICH C18 Silica gel spherically equilibrated with 10 CV of 5% aq. MeCN containing 0.1% trifluoroacetic acid. After washing with a 10 CV of 5% aq. MeCN containing 0.1% trifluoroacetic acid, the peptide was eluted from the column using up to 10 CV of 50% aq. MeCN containing 0.1% trifluoroacetic acid. Fractions containing the eluted peptide were freeze-dried, and the peptide was subsequently dissolved in Tris-HCl (pH=6.5) containing an appropriate amount of NisP leader protease and incubated at 37 °C for 3 h to cleave off the leader peptide. After filtration through a 0.2 μm filter, the core peptide was purified on an Agilent 1260 Infinity HPLC system with a Phenomenex Aeris^™^ C18 column (250 × 4.6 mm, 3.6 μm particle size, 100 Å pore size). Acetonitrile was used as the mobile phase, and a gradient of 15-25% aq. MeCN for thanacin and ripcin and 35-45% aq. MeCN for ripcin B-G over 25 min at 1 mL per min was used for separation. Peptides were eluted at 21-23% MeCN for thanacin and ripcin and 38-41% MeCN for ripcin B-G. After lyophilization, peptides were dissolved in sterilized water and stored in −80°C. The expression levels for designed peptides are listed in Table 1.

### Mass Spectrometry

0.5 μL sample was spotted and dried on the target. Subsequently, 0.5 μL of matrix solution (5 mg/mL α-cyano-4-hydroxycinnamic acid from Sigma-Aldrich dissolved in 50% acetonitrile containing 0.1% trifluoroacetic acid) was spotted on top of the sample. Matrix-assisted laser desorption ionization-time-of-flight (MALDI-TOF) mass spectrometer analysis was performed using a 4800 Plus MALDI TOF/TOF Analyzer (Applied Biosystems) in the linear-positive mode.

### Evaluation of (methyl)Lanthionine Formation

After dissolving the freeze-dried samples in 18 μL of 0.5 M HCL (pH=3), the samples were treated with 2 μL of 100 mg/mL tris[2-carboxyethyl]phosphine in 0.5 M HCL (pH=3) for 30 min at room temperature. Subsequently, 4 μL of 100 mg/mL 1-Cyano-4-dimethylaminopyridinium tetrafluoroborate (CDAP) in 0.5 M HCL (pH=3) was treated to the samples. After incubation at room temperature for 2 hours, the samples were desalted by C-18 ZipTip (Millipore) and analyzed by MALDI-TOF MS ^49^.

### LC-MS/MS Analysis

To get deep insight into the lanthionine bridging pattern we performed LC-MS/MS assay. LC-MS was performed using a Q-Exactive mass spectrometer fitted with an Ultimate 3000 UPLC, an ACQUITY BEH C18 column (2.1 × 50 mm, 1.7 μm particle size, 200 Å; Waters), a HESI ion source and a Orbitrap detector. A gradient of 5-90% MeCN with 0.1% formic acid (v/v) at a flow rate of 0.35 mL/min over 60 min was used. MS/MS was performed in a separate run in PRM mode, selecting the doubly and triply charged ion of the compound of interest.

### Minimum Inhibitory Concentration (MIC) Assay

MIC values were determined by broth micro-dilution, according to the standard guidelines ^40^. Briefly, the test medium was cation-adjusted Mueller-Hinton broth (MHB). Cell concentration was adjusted to approximately 5×10^5^ cells per ml. After 20 h of incubation at 37 °C, the MIC was defined as the lowest concentration of antibiotic with no visible growth. Each experiment was performed in triplicate.

For the nisin resistance protein-producing strain *L. lactis* NZ9000 (pNZ-SV-SaNSR, Em^r^) ^45^ and non-NSR-producing strain *L. lactis* NZ9000 (pNZ8048-Em^r^, Em^r^) ^46^, the MIC tests were performed in GM17, supplemented with erythromycin (5 μg/ml) and nisin (1ng/mL, for maintenance of the induction of NSR). *L. lactis* NZ9000 strains were induced by nisin at a concentration of 5 ng/mL for 3 h before exposing them to antibiotics.

### Time-killing Assay

This assay was performed according to a previously described procedure ^50^. An overnight culture of either *Staphylococcus aureus* ATCC15975 (MRSA) or *Acinetobacter baumannii* LMG01041 was diluted 50-fold in MHB and incubated at 37 °C with aeration at 220 r.p.m. Bacteria were grown to an OD_600_ of 0.6, and then the concentration of cells was adjusted to ≈5×10^6^ cells per mL for both strains. Bacteria were then challenged with 10×MIC antimicrobials in glass culture tubes at 37 °C and 220 r.p.m. Bacteria not treated with peptides were used as a negative control. At desired time points, two hundred μl aliquots were taken, centrifuged at 8,000 g for 2 min, and resuspended in 200 μl of MHB. Ten-fold serially diluted samples were plated on MHA plates. After incubation at 37 °C overnight, colonies were counted and the c.f.u. per ml was calculated. Each experiment was performed in triplicate.

### Fluorescence Microscopy Assay

*Staphylococcus aureus* ATCC15975 (MRSA) or *Acinetobacter baumannii* LMG01041 was grown to an OD_600_ of 0.8. The culture was pelleted at 4,000g for 5 min and washed three times in MHB. After normalization of the cell density to an OD_600_ of 0.2 in MHB, a two-fold MIC value concentration of antimicrobials was added. Simultaneously, SYTO^®^9 and propidium iodide (LIVE/DEAD Baclight Bacterial Viability Kit, Invitrogen) were added to the above cell suspensions. After incubation at room temperature for 15 min, peptides were removed and washed three times with MHB. Then the cell suspensions were loaded on 1.5 % agarose pads and analyzed by DeltaVision Elite microscope (Applied precision).

### Bacterial Growth Assay to Monitor Lipid II and Lipopolysaccharide (LPS) Binding of Antibiotics

Briefly, the test medium was cation-adjusted Mueller-Hinton broth (MHB). The cell concentration was adjusted to approximately 5×10^5^ cells per ml, and then cells were transferred to a 96-well plate and thereafter challenged with antibiotics at a concentration of 2×MIC either with or without lipid II ^51^ (10 μM; provided by Prof. Eefjan Breukink)/LPS (100 μg/mL; Merck, L2880-100MG). The absorbance values were measured by using a Varioskan^™^ LUX multimode microplate reader (Thermo Fisher Scientific) at 600 nm. Each experiment was performed in triplicate.

## ASSOCIATED CONTENT

### Supporting Information

The Supporting Information is available free of charge at Critical area of plasmids encoding the designed peptide genes; MALDI-TOF MS data of a free Cys containing peptide before (blue) and after (red) treatment with CDAP; Full LC-MS/MS spectrums of thanacin and ripcin; Expression of peptides measured by SDS-tricine gel (From TCA precipitation); Digestion of nisin by chymotrypsin for production of nisin(1-20); MALDI-TOF MS data of HPLC-purified nisin(1-20); Strain and plasmids used in this study; Primers for PCRs used in this study.

## AUTHOR INFORMATION

**Corresponding Author** *

### Author contributions

O.P.K. and X.Z. conceived the project and strategy. O.P.K. supervised and corrected the manuscript. X.Z. designed and carried out the experiments, analyzed data, and wrote the manuscript. All authors contributed to and commented on the manuscript text and approved its final version.

## ACKNOWLEDGMENTS

X.Z. was financially supported by the Netherlands Organization for Scientific Research (NWO), research program TTW (Project No. 17241). We thank Prof. Eefjan Breukink (Membrane Biochemistry and Biophysics, Department of Chemistry, Faculty of Science, Utrecht University, Utrecht, The Netherlands) for providing us the purified lipid II. We thank Marcel P. de Vries for support with LC-MS/MS.

