## Supplemental Figures and Tables for "Nisin- and ripcin-derived hybrid lanthipeptides display selective antimicrobial activity against *Staphylococcus aureus*"

A

B

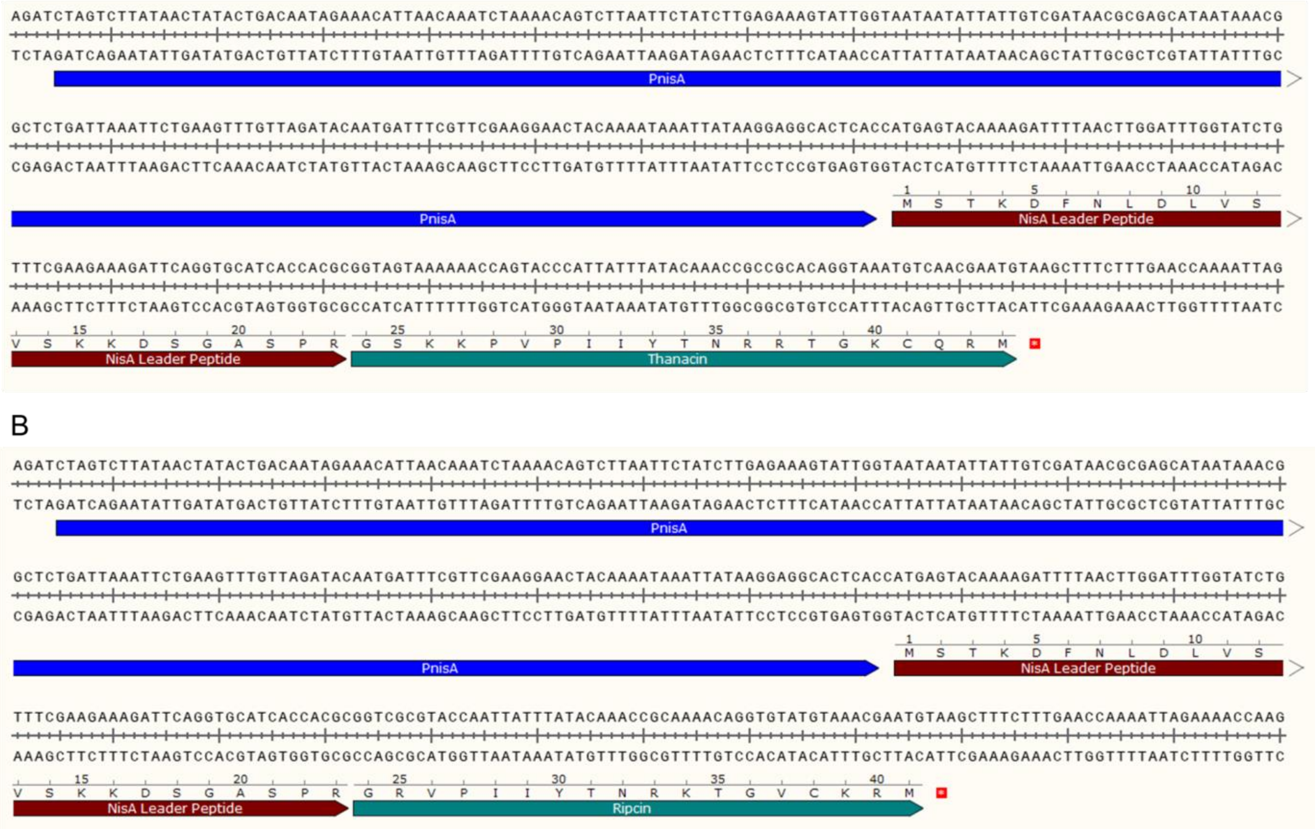

**Figure S1** Critical area of plasmids encoding the designed peptide genes. A, part of the sequence of the plasmid encoding thanacin. B, part of the sequence of the plasmid encoding ripcin.

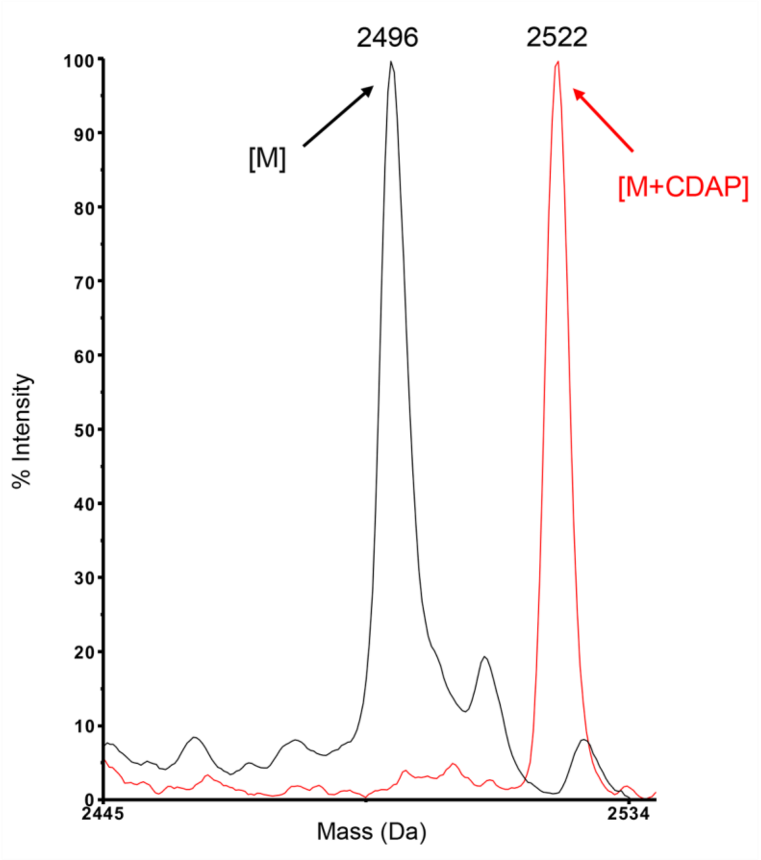


**Figure S2** MALDI-TOF MS data of a free Cys containing peptide before (blue) and after (red) treatment with CDAP.

A

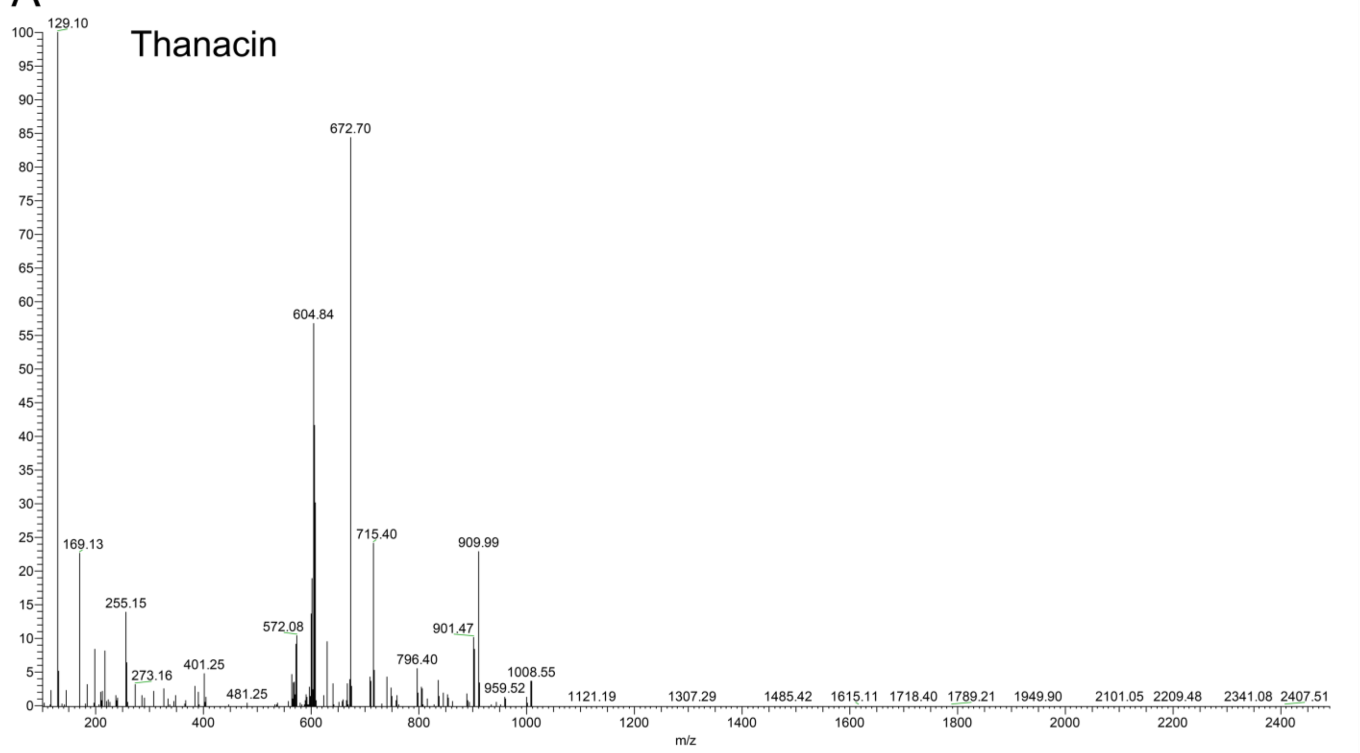


B

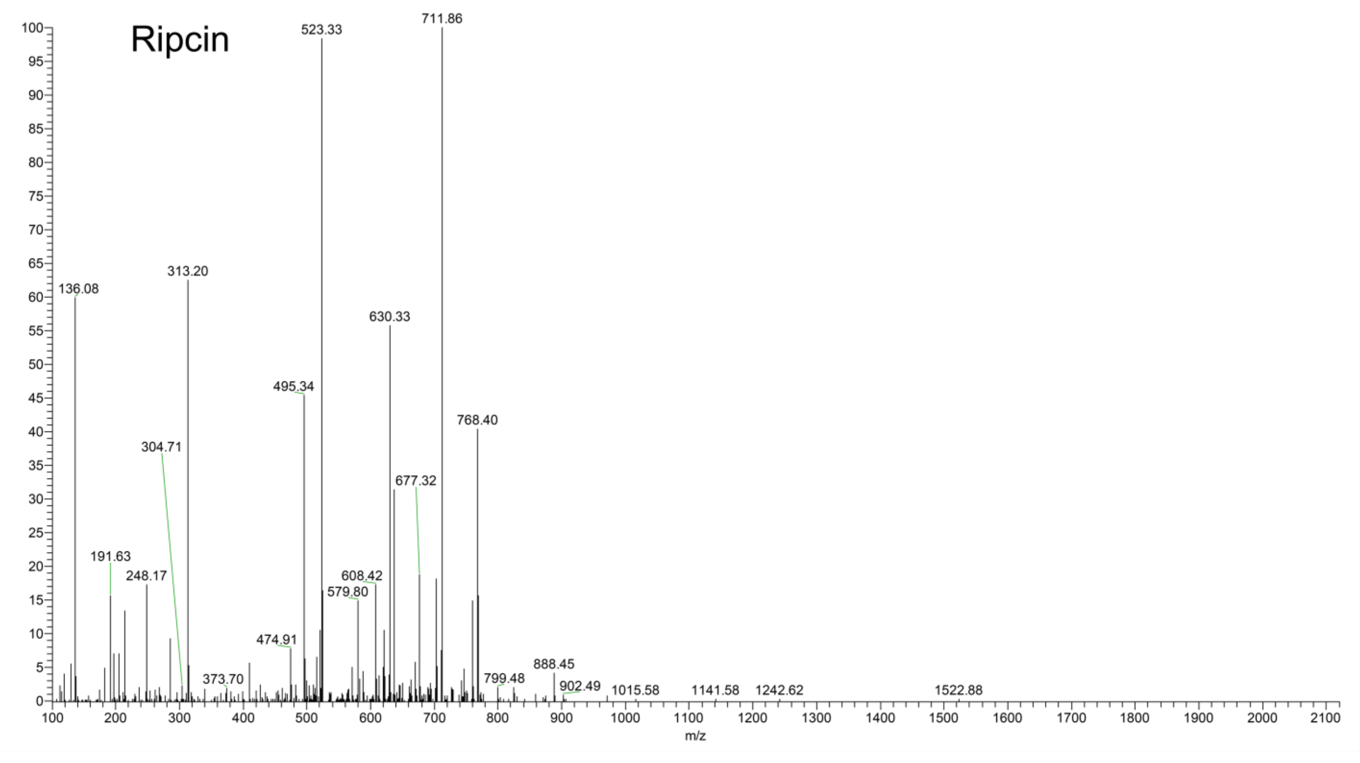


**Figure S3** Full LC-MS/MS spectrums of thanacin and ripcin.

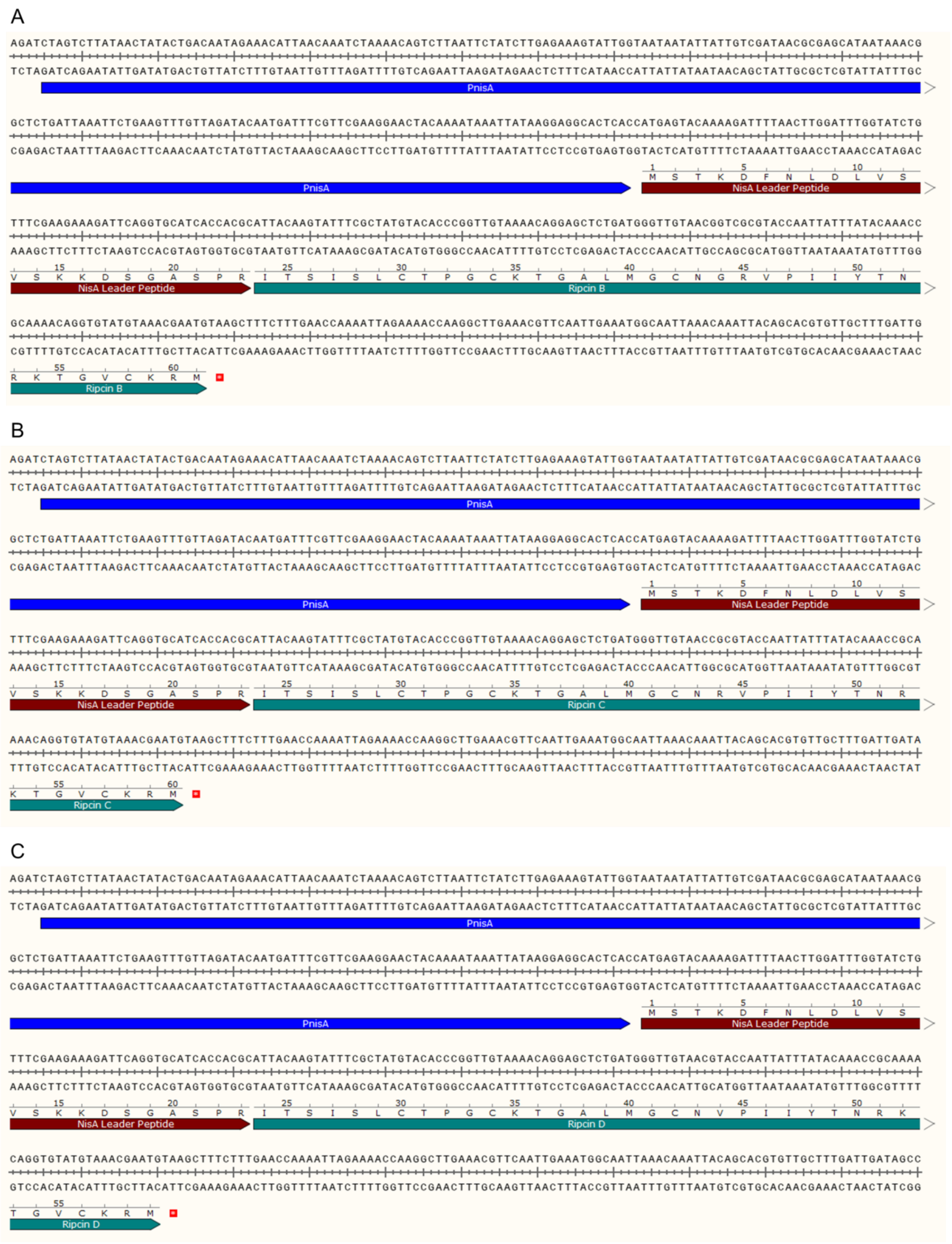

**Figure S4** Critical area of plasmids encoding the designed peptide genes. A, part of the sequence of the plasmid encoding ripcin B. B, part of the sequence of plasmid encoding ripcin C. C, part of the sequence of the plasmid encoding ripcin D.

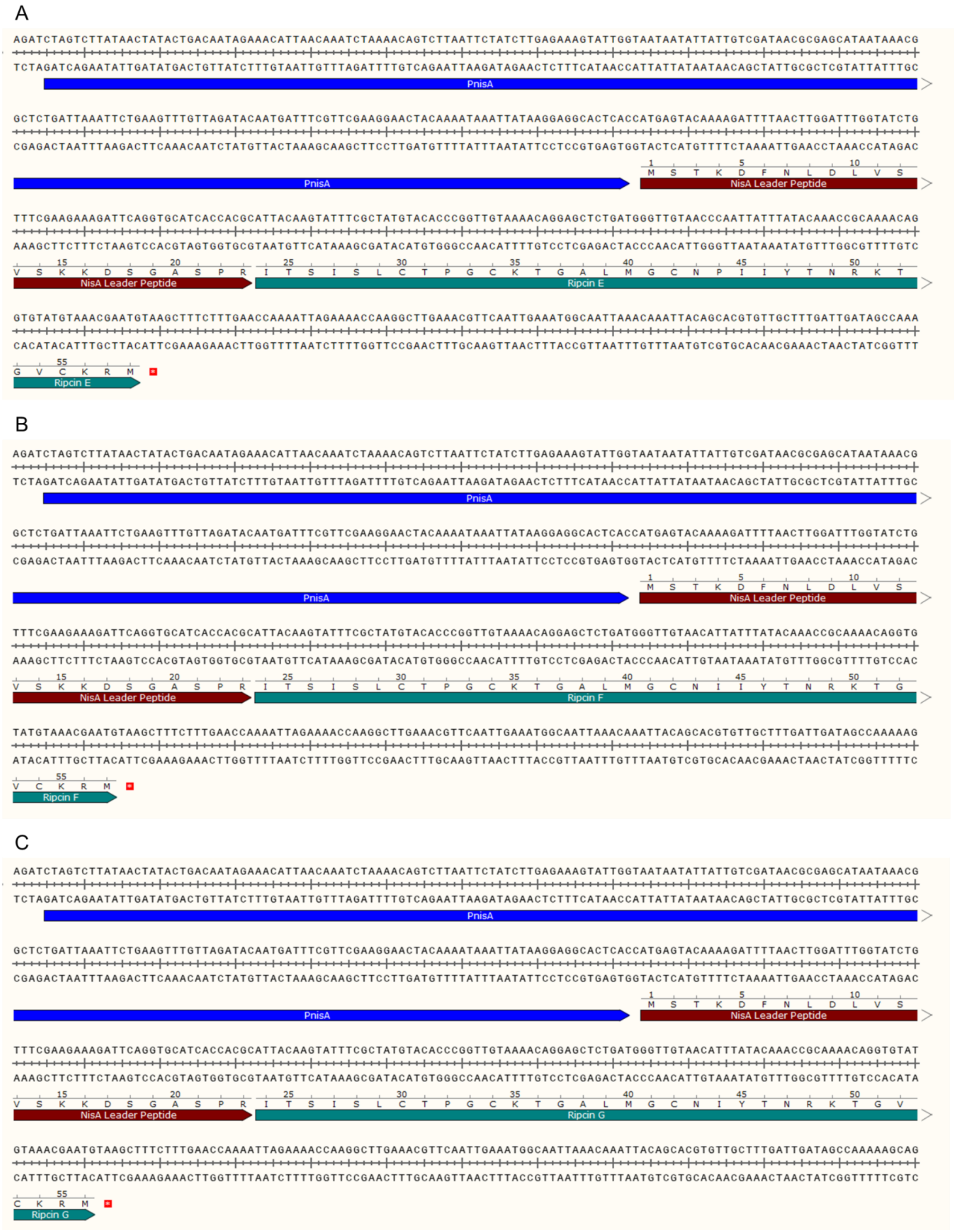

**Figure S5** Critical area of plasmids encoding the designed peptide genes. A, part of the sequence of plasmid encoding ripcin E. B, part of the sequence of plasmid encoding ripcin F. C, part of the sequence of plasmid encoding ripcin G.

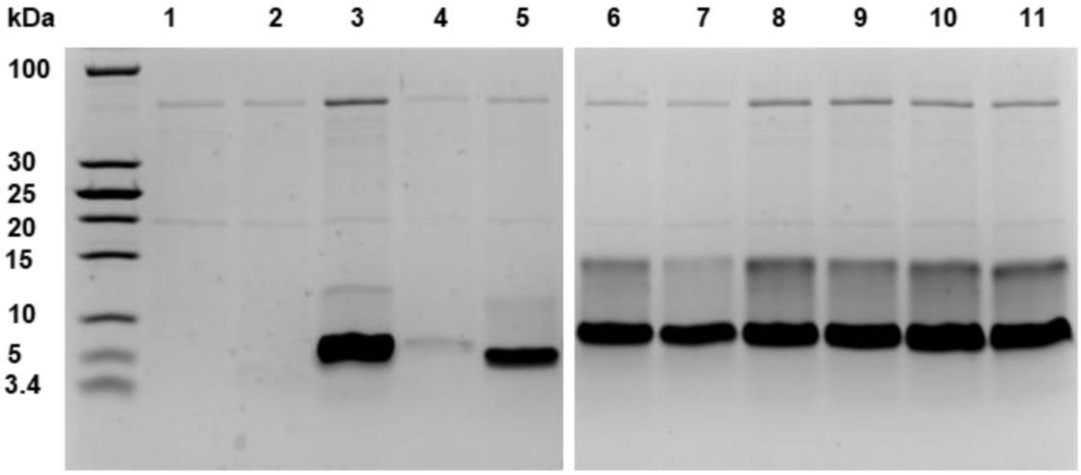


**Figure S6** Expression of peptides measured by SDS-tricine gel (From TCA precipitation), lane 1: pNZ8048, lane 2: pNZ8048 with pIL3BTC, lane 3: Nisin (1-22), lane 4: Thanacin, lane 5: Ripcin, lane 6: Ripcin B, lane 7: Ripcin C, lane 8: Ripcin D, lane 9: Ripcin E, lane 10: Ripcin F, lane 11: Ripcin G.

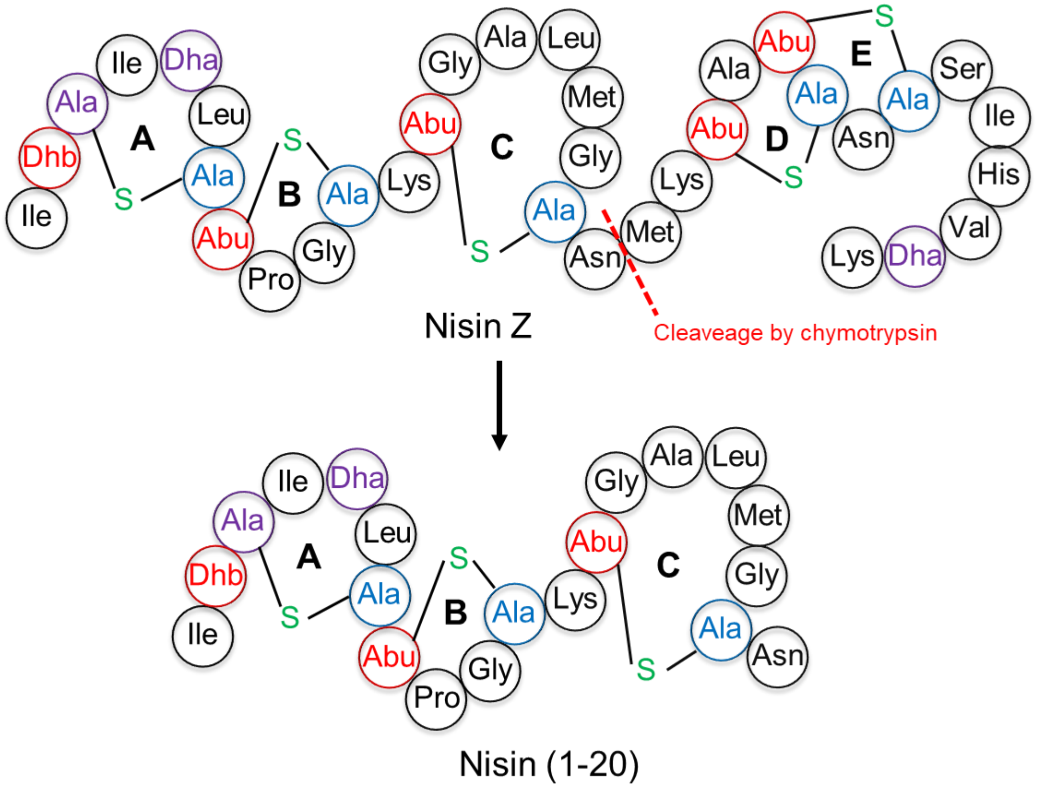


**Figure S7** Digestion of nisin by chymotrypsin for production of nisin(1-20). Dha: dehydroalanine, Dhb: dehydrobutyrine, Abu: aminobutyric acid.

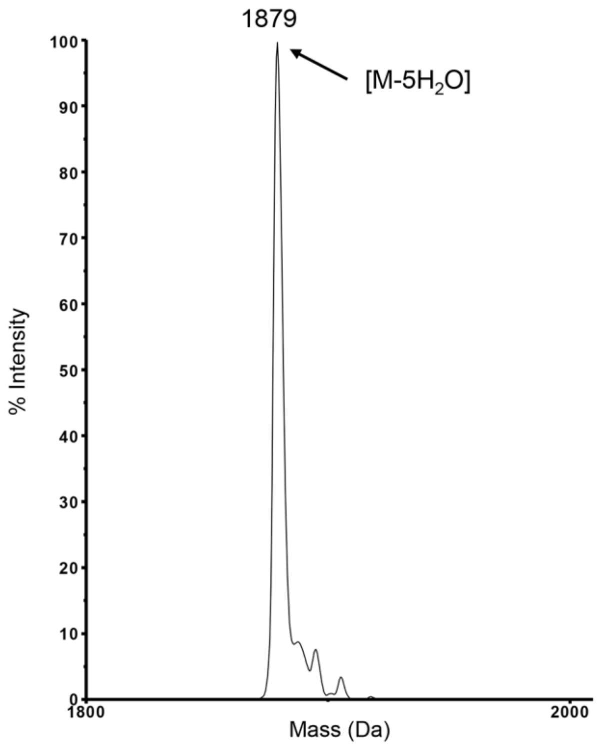


**Figure S8** MALDI-TOF MS data of HPLC-purified nisin(1-20).

**Supplementary Table 1.** Stains and plasmids used in this study.

| Strains or plasmids | Characteristics and purpose | Source or reference |
|---|---|---|
| *Strains* | | |
| *Lactococcus lactis NZ9000* | MG1363 derivative; NisRK+, plasmid construction, plasmid maintenance and protein expression. | [1] |
| *Lactococcus lactis NZ9000* | MG1363 derivative; NisRK+, pNZ8048-$Em^r$, indicator strain, $Em^r$ | [2] |
| *Lactococcus lactis NZ9000* | MG1363 derivative; NisRK+, pNZ-SV-SaNSR, nisin resistant protease producer, indicator strain, $Em^r$ | [3] |
| *Staphylococcus aureus* | ATCC15975 (MRSA), indicator strain | |
| *Staphylococcus aureus* | LMG10147, indicator strain | |
| *Enterococcus faecium* | LMG16003 (VRE), indicator strain | |
| *Enterococcus faecalis* | LMG16216 (VRE), indicator strain | |
| *Bacillus cereus* | ATCC 14579, indicator strain | |
| *Acinetobacter baumannii* | LMG01041, indicator strain | |
| *Shigella flexneri* | ATCC29903, indicator strain | |
| *Escherichia coli* | ATCC25922, indicator strain | |
| *Klebsiella pneumoniae* | LMG20218, indicator strain | |
| *Pseudomonas aeruginosa* | LMG6395, indicator strain | |
| *Enterobacter cloacae* | LMG02783, indicator strain | |
| *Plasmids* | | |
| pNisin(1-22) | pNZ8048 derivative, template for plasmid construction, expression NisA leader-Nisin(1-22), $Cm^r$ | [4] |
| pThanacin | pNZ8048 derivative, expression NisA leader-Thanacin, $Cm^r$ | This work |
| pRipcin | pNZ8048 derivative, expression NisA leader-Ripcin, $Cm^r$ | This work |
| pRipcin B | pNZ8048 derivative, expression NisA leader-Ripcin B, $Cm^r$ | This work |
| pRipcin C | pNZ8048 derivative, expression NisA leader-Ripcin C, $Cm^r$ | This work |
| pRipcin D | pNZ8048 derivative, expression NisA leader-Ripcin D, $Cm^r$ | This work |
| pRipcin E | pNZ8048 derivative, expression NisA leader-Ripcin E, $Cm^r$ | This work |
| pRipcin F | pNZ8048 derivative, expression NisA leader-Ripcin F, $Cm^r$ | This work |
| pRipcin G | pNZ8048 derivative, expression NisA leader-Ripcin G, $Cm^r$ | This work |
| pIL3-BTC | $P_{nis}$ nisBTC, expression nisBTC enzyme, $Em^r$ | [5] |

**Supplementary Table 2.** Primers for PCRs used in this study.

| New plasmids | Templates | Primers | Nucleic acid sequences (5' to 3') | Characteristics (5'-lable) |
|---|---|---|---|---|
| pThanacin | pNisin(1-22) | PM1 | CGCACAGGTAAATGTCAACGAATGTAAGCTTTCTTTGAACCAAAATTAGAAAACCAAG | 5'- phosphorylation |
| | | PM2 | GCGGTTTGTATAAATAATGGGTACTGGTTTTTTACTACCGCGTGGTGATGCACCTGAATC | |
| pRipcin | pNisin(1-22) | PM3 | GTGTATGTAAACGAATGTAAGCTTTCTTTGAACCAAAATTAGAAAACCAAG | 5'- phosphorylation |
| | | PM4 | CTGTTTTGCGGTTTGTATAAATAATTGGTACGCGACCGCGTGGTGATGCACCTGAATC | |
| pRipcin B | pNisin(1-22) | PM3 | GTGTATGTAAACGAATGTAAGCTTTCTTTGAACCAAAATTAGAAAACCAAG | 5'- phosphorylation |
| | | PM5 | CTGTTTTGCGGTTTGTATAAATAATTGGTACGCGACCGTTACAACCCATCAGAGCTCCTGTTTTAC | |
| pRipcin C | pNisin(1-22) | PM3 | GTGTATGTAAACGAATGTAAGCTTTCTTTGAACCAAAATTAGAAAACCAAG | 5'- phosphorylation |
| | | PM6 | CTGTTTTGCGGTTTGTATAAATAATTGGTACGCGGTTACAACCCATCAGAGCTCCTGTTTTAC | |
| pRipcin D | pNisin(1-22) | PM3 | GTGTATGTAAACGAATGTAAGCTTTCTTTGAACCAAAATTAGAAAACCAAG | 5'- phosphorylation |
| | | PM7 | CTGTTTTGCGGTTTGTATAAATAATTGGTACGTTACAACCCATCAGAGCTCCTGTTTTAC | |
| pRipcin E | pNisin(1-22) | PM3 | GTGTATGTAAACGAATGTAAGCTTTCTTTGAACCAAAATTAGAAAACCAAG | 5'- phosphorylation |
| | | PM8 | CTGTTTTGCGGTTTGTATAAATAATTGGGTTACAACCCATCAGAGCTCCTGTTTTAC | |
| pRipcin F | pNisin(1-22) | PM3 | GTGTATGTAAACGAATGTAAGCTTTCTTTGAACCAAAATTAGAAAACCAAG | 5'- phosphorylation |
| | | PM9 | CTGTTTTGCGGTTTGTATAAATAATGTTACAACCCATCAGAGCTCCTGTTTTAC | |
| pRipcin G | pNisin(1-22) | PM3 | GTGTATGTAAACGAATGTAAGCTTTCTTTGAACCAAAATTAGAAAACCAAG | 5'- phosphorylation |
| | | PM10 | CTGTTTTGCGGTTTGTATAAATGTTACAACCCATCAGAGCTCCTGTTTTAC | |
| Sequencing primer PrXZ12 | | | CTATCAATCAAAGCAACACGTGC | |
